# Genus-wide sequencing supports a two-locus model for sex-determination in *Phoenix*

**DOI:** 10.1101/245514

**Authors:** Maria F. Torres, Lisa S. Mathew, Ikhlak Ahmed, Iman K. Al-Azwani, Robert Krueger, Diego Rivera, Yasmin A. Mohamoud, Andrew G. Clark, Karsten Suhre, Joel A. Malek

**Author notes:** Corresponding Author: Joel A. Malek. Current address: Department of Biological Sciences, University of Cincinnati, Cincinnati, OH.

## Abstract

The date palm tree is a commercially important member of the genus *Phoenix* whose 14 species are all dioecious with separate male and female individuals. Previous studies identified a multi-megabase region of the date palm genome linked to sex and showed that dioecy likely developed in *Phoenix* prior to speciation. To identify genes critical to sex determination we sequenced the genomes of 28 *Phoenix* trees representing all 14 species. Male-specific sequences were identified and extended using phased single molecule sequencing or BAC clones to distinguish X and Y alleles.

Here we show that only four genes contain sequences conserved in all analyzed males, likely identifying the changes foundational to dioecy in *Phoenix*. The majority of these sequences show similarity to a single genomic locus in the closely related oil palm. *CYP703* and *GPAT3*, two genes critical to male flower development in other monocots, appear fully deleted in females while maintained as single copy in males. A LOG-like gene appears translocated into the Y chromosome and a cytidine deaminase-like appears at the border of a chromosomal rearrangement. Our data supports a two-mutation model for the evolution from hermaphroditism to dioecy through a gynodioecious intermediate.

## Introduction

The origin and evolution of separate sexes is a subject that has intrigued scientists for many years. The presence of separate male and female individuals (dioecy) ensures certain advantages such as outcrossing and improved fertility by allocation of resources^1^. One model that describes the transition from cosexuality to dioecy involves two mutations in a pair of autosomes: a recessive male sterility mutation in the proto-X chromosome that creates females and a dominant female sterility mutation in the proto-Y chromosome that creates males with recombination ultimately suppressed between them^2,3^. Lack of recombination in the heterogametic sex (XY) promotes the accumulation of mutations and repetitive elements, which in turn leads to chromosome degeneration (Reviewed in^4^). While most species of animals have separate sexes, only 5–6% of all plant species are dioecious^5^. Various genomic approaches^6^ have allowed the identification of sex-associated genes in papaya (*Carica papaya*), white campion (*Silene latifolia*), grapevine (*Vitis vinifera*), poplars (*Populus sp*.) and the herb *Mercurialis annua*^7^–11. However, information about how these genes control plant sex remains elusive save for few notable exceptions for melon and persimmon ^12,13^.

The dioecious date palm (*Phoenix dactylifera*) has recently emerged as a system to study plant sex chromosomes. The female date palm produces the commercially important dates and is of high agricultural value in North Africa, the Middle East and South East Asia. We previously genetically mapped a sex-linked region to the long arm of linkage group 12 and estimated this region to span 13 Mb, or 2% of the genome^14,15^. While sex-linked markers and genes have been identified in date palm, sex-determination genes remain to be identified^14,16^–^18^. Phylogenetic analysis of one of these sex-linked genes suggests that dioecy preceded speciation in the genus *Phoenix*^18^. All fourteen members of the genus *Phoenix* are dioecious, while related genera in the palm family are not^19^. Furthermore, hybridization can occur among species with fertile progeny in some cases. Hypothesizing a single origin of dioecy, we designed a study that combines *de novo* whole-genome sequencing and comparative genomics across all fourteen members of *Phoenix* with the aim to identify the original genes responsible for sex determination in the genus. We report the identification of four candidate sex-determination genes, three of them completely absent in all female *Phoenix* individuals, but present in two closely related hermaphroditic palms. We discuss the putative role of these genes and propose a model for the origination of dioecy in *Phoenix*.

## Results

### Identification of Male-Specific Sequences

Identification of sequences (16 bp kmers) unique to either males or females in each species (Table 1 and Supplementary Table 1) revealed that only kmers specific to males existed in all 14 species (Figure 1a). Indeed, 1653 kmers (Supplementary Table 2) were shown to be present in all 13 males and absent in all 14 females tested (a male from *P. pusilla* could not be identified for sequencing). In contrast, kmers present in females and absent in males were not present in more than 8 species of the genus. This was expected in a genus that employs an XY sex determination system with the male being the heterogametic sex.

**Figure 1.**
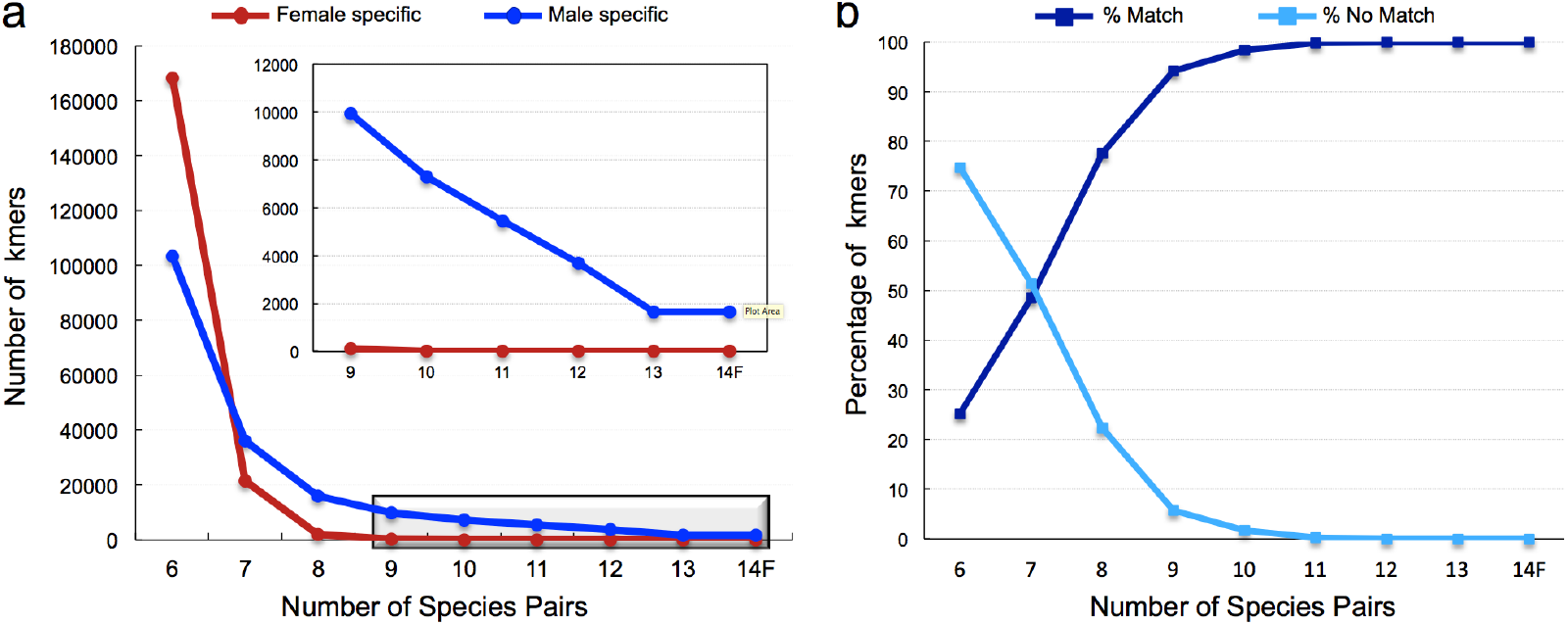
Identification of male-specific kmers in the genus *Phoenix*. (a)16 bp kmers identified only in the male or female individual for each species were tested for their presence in other species. As expected for an XY sex determination system, kmers specific to males maintained counts through all species tests while female-specific kmers disappeared after requiring presence in more than 8 species. (b) Kmers specific to male *Phoenix* were enumerated against the male-specific BAC contigs. Kmers specific to males in at least 12 species are fully covered by the sequenced BACs in this study.

**Table 1.**
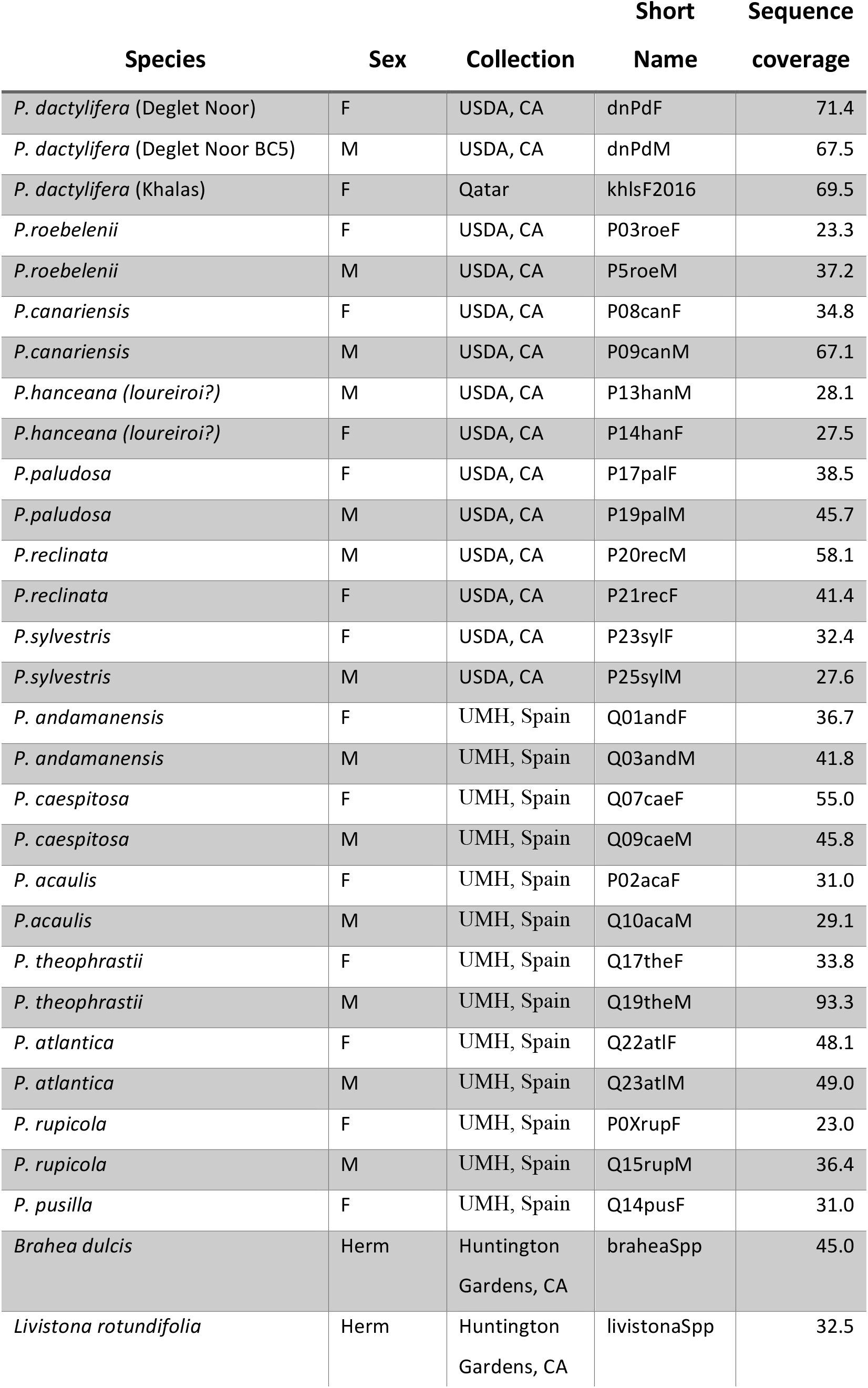
Genomes sequenced in this study.

A date palm male BAC library was probed to identify clones harboring the kmer sequences. Where possible, a BAC clone representing both the X and Y allele of a locus was sequenced. Additionally, phased single molecule sequencing (10X Genomics) scaffolds of the same male genome were searched for male-specific kmers. This resulted in 11 BAC contigs and 30 phased sequence scaffolds spanning ~5.5 Mb (~11 Mb of male and female alleles) in the region of interest (Supplementary Table 3). These BAC contigs and sequence scaffolds covered 100% of the male-specific kmers present in at least 12 or more species (Figure 1b).

Annotation of contigs harboring the highest density of kmers specific to all males in the genus (Figure 2) revealed the presence of just a few genes surrounded by highly repetitive sequence. These included full-length genes with similarity to *CYP703, GPAT3, LOG* and, to a lesser extent, cytidine deaminase for which there was a male and female allele. The BAC contig containing cytidine deaminase also included a degraded copy of the MAF1-like gene that contained no male-specific kmers.

**Figure 2.**
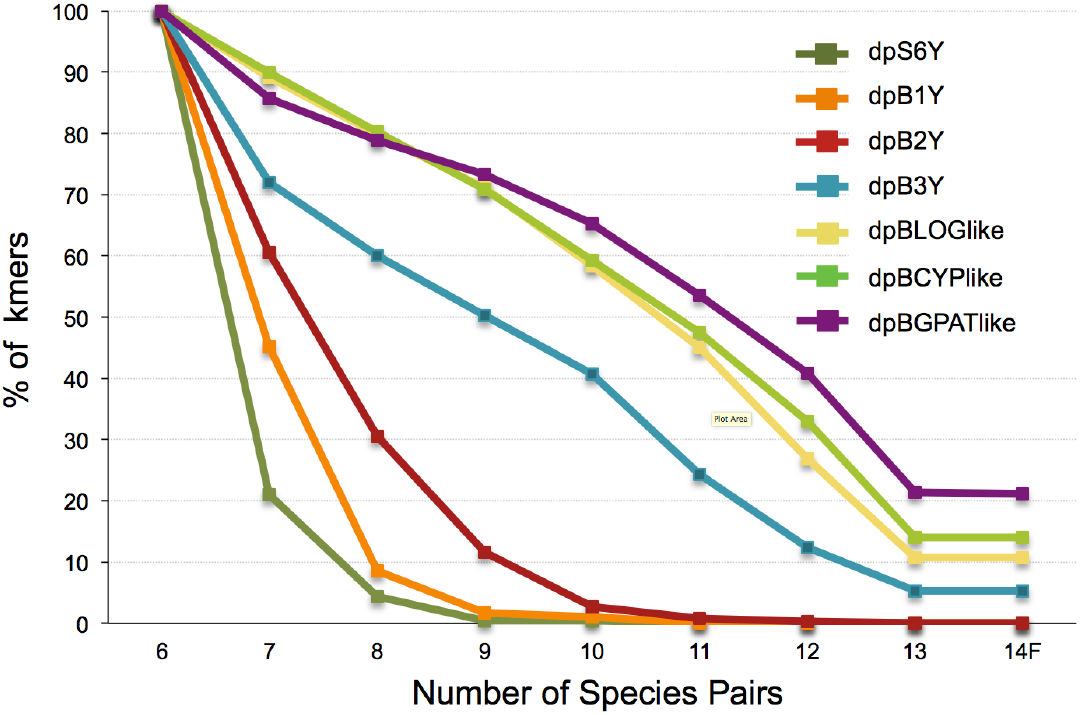
Male-specific BAC sequences contain varying numbers of male-specific kmers. Percent of kmers matching in 6 species were used to normalize for size of the BAC contig. The GPAT3 contig maintained the largest percentage of kmers through the 14 species suggesting possibly stronger selection on this gene or a more recent loss of recombination. It was followed by CYP703, LOG and cytidine deaminase (dpB3Y contig) gene. For comparison see synonymous mutation rates in the same genes.

### Analysis of male-specific genes

Searches of the NCBI database indicated that one of the male-specific genes is a Cytochrome P450 (CYP) similar to *CYP703A* (LOC105059962) in oil palm (Supplementary Figure 1). We performed a phylogenetic analysis of the predicted CYP-like proteins from *Phoenix, Brahea* and *Livistona* with 32 CYP450 proteins from 11 plant species^20,21^. Our results indicate that the CYP gene identified in this study belongs to the CYP703 family and is a putative ortholog of *CYP703A3* from rice (not shown). We found no evidence of complete or partial sequences of *CYP703* in the published female reference genomes or the transcriptome data available from date palm^14,22^.

The second identified male-specific gene is highly similar to a glycerol-3-phosphate acyltransferase 6-like (GPAT6-like) gene (LOC105059961) in oil palm. Phylogenetic analysis was conducted on 24 GPAT genes including the GPAT-like gene from 3 species identified in this study, the oil palm GPAT6-like, the eight characterized *AtGPAT* genes from Arabidopsis^23^, and the twelve *OsGPAT*-like genes from rice available in the NCBI database. Our results indicate that the GPAT-like gene from the four palms belong to the GPAT 1/2/3 (annotated GPAT3) clade^24^, considered a more diverse clade than GPAT4/6/8 or GPAT 5/7 clades (Supplementary Figure 2).

The third highly conserved male-specific gene is similar to a family of genes termed *Lonely-Guy* (LOG). We performed a phylogenetic analysis of all the date palm and oil palm *LOG* genes available in the NCBI database and included the rice *LOG* genes, some of which have been previously identified as crucial for normal shoot and floral meristem development^25^. Our phylogenetic analysis suggested that the LOG-like gene is a paralog of the date palm autosomal gene Pd_LOC103701078 (Supplementary Figures 3 and 4). Furthermore, this autosomal gene appears to be the ortholog of LOC105055182, a gene found in chromosome 12 in oil palm as indicated by synteny across loci. This suggests that the male-specific *LOG* gene is absent from oil palm and is likely the result of a duplication of the autosomal gene that occurred after divergence of *Elaeis* and *Phoenix*. We identified male-specific LOG kmers in the hermaphroditic palms *Brahea* and *Livistona*, suggesting that this duplication occurred in a common ancestor of the subfamily Coryphoideae. Analysis also indicated that the LOG-like gene is related to the *OsLOG5* and *OsLOG9* genes in rice.

**Figure 3.**
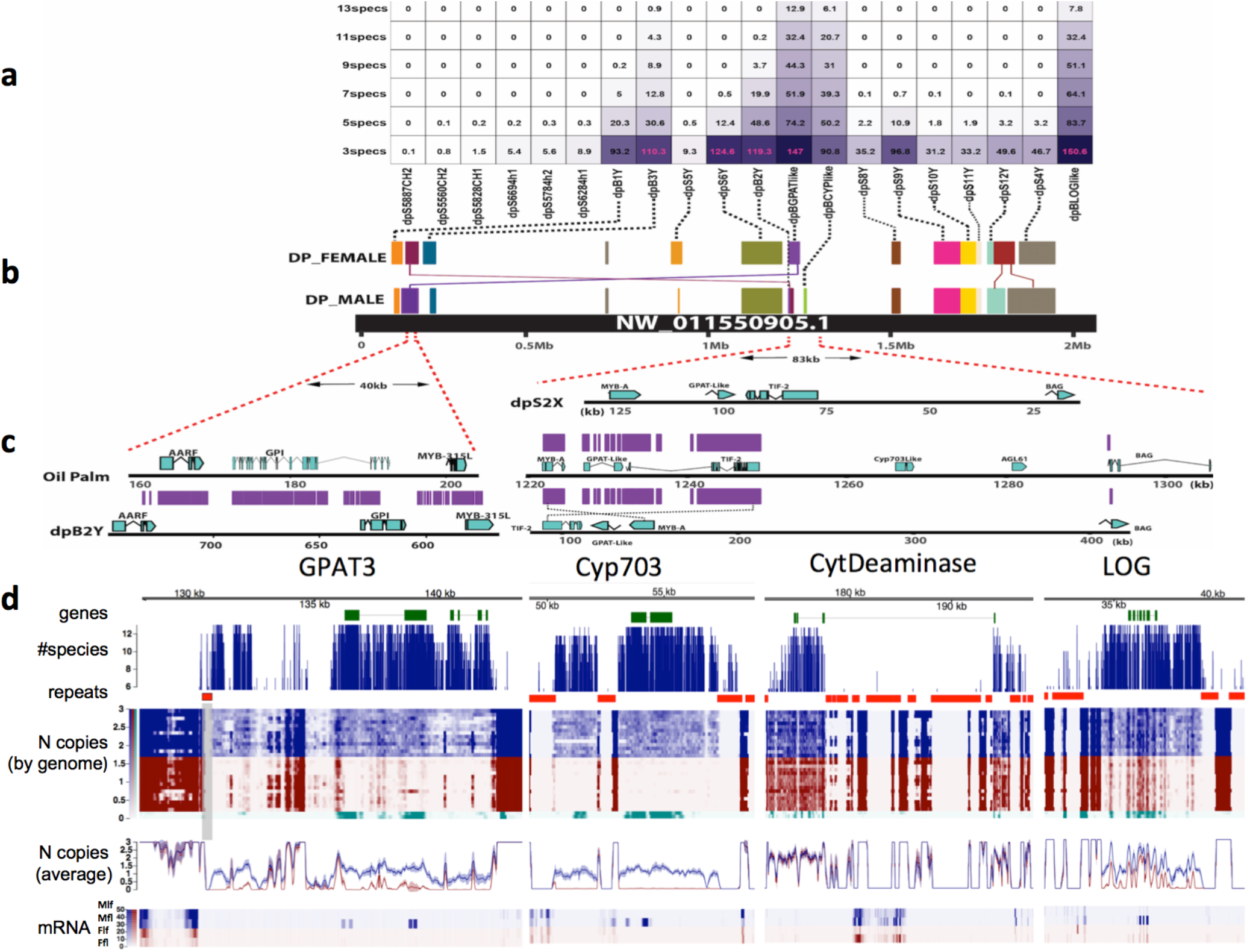
The sex determination region in *Phoenix*. (a) *Phoenix* male-specific kmers/bp in date palm scaffolds based on number of species compared. Scaffolds with male-specific kmers had similarity to a single locus in oil palm. The only gene not deriving from the region was LOG-like. (b) Sequencing of representatives of both male and female alleles in date palm showed coverage of significant sections of the oil palm scaffold NW_011550905.1. (c) Comparison of the X and Y alleles to the orthologous region revealed a deletion of the *GPAT3* and *CYP703* gene in the X chromosome. A similar *GPAT3* deletion is found in the Y chromosome although the *GPAT3* gene does exist elsewhere. The Y region could not be extended further to ensure the *CYP703* was deleted from this region and may indeed reside downstream of *TIF-2*, maintaining original gene order with expanded repeats surrounding it. In oil palm, the cytidine deaminase (not shown) resides just downstream of the MYB315L. The region showed a fusion/inversion with respect to the oil palm locus joining the region downstream of MYB315L to the region upstream of BAG-like in date palm. (d) Analysis of male-specific kmers present in all species of *Phoenix* identified only 4 scaffolds dpBGPATlike containing *GPAT3*, dpBCYPlike containing *CYP703*, dpB3Y containing cytidine deaminase like and dpBLOGlike containing *LOG*. The most widely conserved kmers were focused in gene and apparently gene regulatory regions (# species track). DNA sequence coverage of each genome was normalized to 2*N* and plotted as a heatmap. Only males showed coverage in these regions (*N* copies, by genome track). Blue rows represent males, red rows represent females from the various *Phoenix* species. Green rows represent hermaphrodite palms *Brahea* and *Livistona*. The average coverage for all males indicated these regions were likely at 1*N* except for the LOG gene that has an autosomal paralog (*N* copies, average track). Gene expression (mRNA track) showed that *GPAT3, CYP703* and *LOG* were all expressed only in male flower (Mfl) and was either very low or undetectable in male leaf (Mlf), female flower (Ffl) and female leaf (Flf).

**Figure 4.**
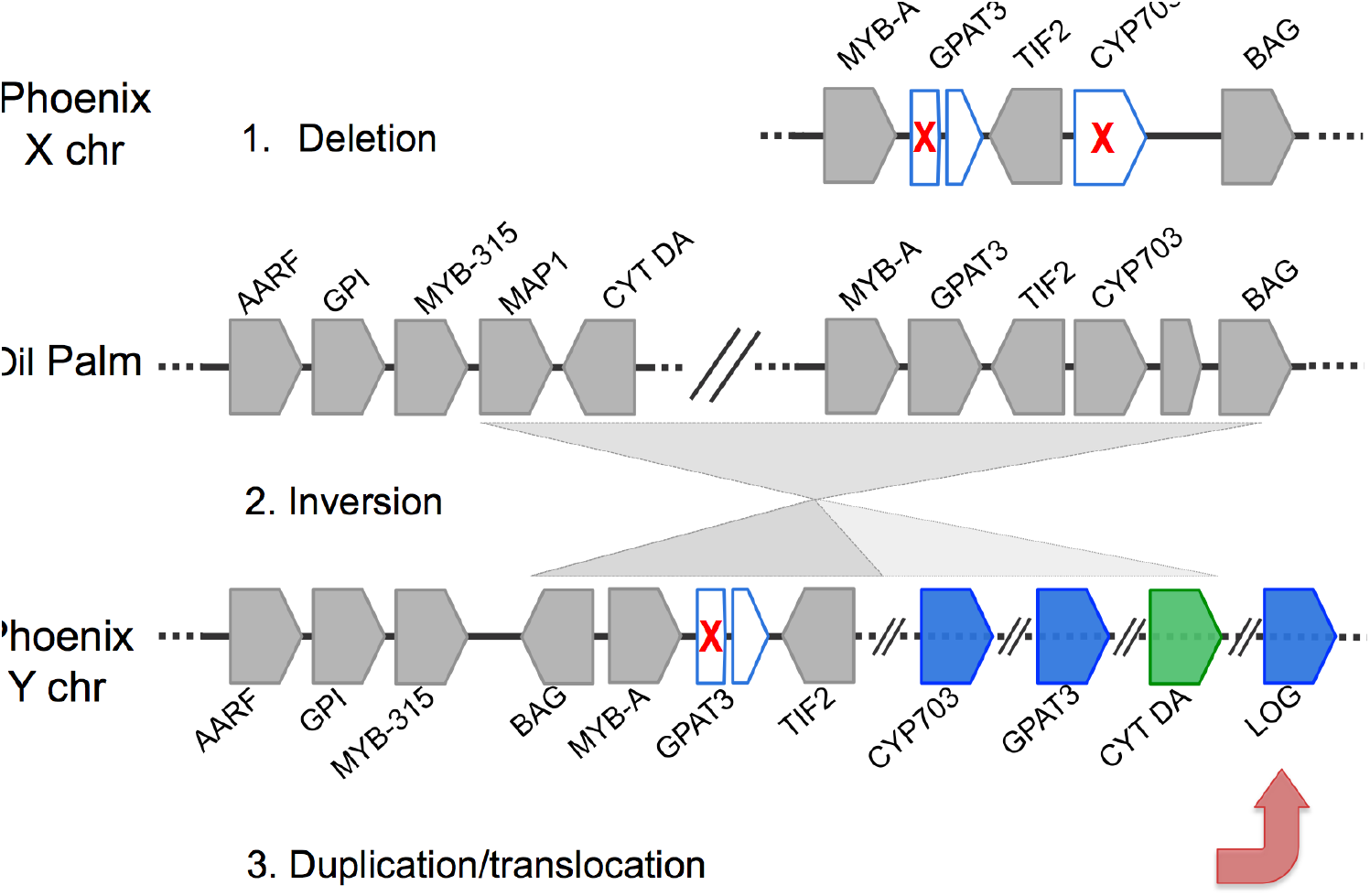
A model for the development of dioecy in the genus *Phoenix*. The genes in the region as found in female scaffold dpS2X, oil palm scaffold NW_011550905.1, and male contig dpB2Y. The fact that the male-specific sequences are focused in this region suggests it may be the origin of sex determination in *Phoenix*. The first step would be the deletion of *GPAT3* and/or *CYP703*, two genes known to be critical to male flower formation, leading to gynodioecy and formation of a proto-X chromosome. An inversion of the corresponding region on the normal chromosome moving the CYT DA gene would create a proto-Y chromosome with recombination arrest. This would be followed by a duplication and translocation into the region of the *LOG* gene creating the final Y chromosome with suppression of female flower development. NCBI GENE IDs as follows AARF: LOC105059738, GPI: LOC105059739, MY315L: LOC105059740, MAP-1: LOC105059742, CYT DA: LOC105059743, MYB-A: LOC105059783, GPAT3: LOC105059961, GPAT3tr: truncated GPAT3, TIF2: LOC105059784, CYP703: LOC105059962, BAG: LOC105059785. The intervening gene between CYP703 and BAG was not present in either male or female alleles of date palm.

Cytidine deaminase was the only gene with genus-wide male-specific kmers for which an X and Y copy was identified. Phylogenetic analysis of this gene indicated that, except for *P. rupicola*, the male and female alleles clustered together suggesting that both alleles evolved before speciation (Supplementary Figure 5). Upon investigation, it was noted that *P. rupicola* had large numbers of male-specific kmers surrounding exonic regions, but did not have any within the exons used here for multiple sequence alignment. Only the X and Y alleles from *P. dactylifera, P. atlantica* and *P.theophrasti* were clustered with high branch support values.

Analysis of the synonymous mutation rate (dS) in the four genes for all *Phoenix* species compared to the ortholog in oil palm revealed different divergence rates among the genes. Across all species, *CYP703* had a mean dS of 0.30 with standard deviation (S.D.) of 0.01. and cytidine-deaminase had a similar rate with a mean dS of 0.28 and S.D. of 0.02 These were followed by *LOG* (mean dS 0.21, S.D. 0.01) and *GPAT3* (mean dS 0.20, S.D. 0.01). In comparison, a set of 22 genes conserved in all plant genomes termed “universal single copy orthologs”^26^ showed a mean dS of 0.14 and S.D. 0.03. Possible reasons for the accelerated rate of synonymous substitution in the sex-specific genes are explored in the Discussion.

Using the alignments of the sequences of the four genes across the *Phoenix* species and the two monoecious palm species, we could infer the relative time to most recent common ancestry of each gene. First we note that all four genes have a gene genealogy that is concordant with a monophyletic origin across the date palm species. Using the TimeTree approach^27^ we inferred the proportion of the time back to common ancestry of the monoecious and dioecious palms that each gene tree converged. For CYP703, GPAT3 and LOG, these values were 13.4%, 15.2% and 9.2%, consistent with their arrival on the Y chromosome. Cytidine deaminase could not be analyzed this way because the monoecious allele falls within the date palm clade, suggesting there might have been some X-Y gene conversion or other complication.

### Inference of natural selection

Alignments of each of the four genes provide a chance to assess whether divergence at synonymous and nonsynonymous sites occurs among the Y-linked genes in the genus *Phoenix* according to neutral divergence. CYP703 had to retain function on the Y chromosome in order to retain fertility of male flowers (discussed below), and it displays a value of ω (*dN*/*dS*) of 0.16. In addition, codon-based tests in HyPhy^28^ revealed many individual codons showing strong purifying selection, and many pairwise contrasts among species also consistently showed a strong signal of purifying selection. One might make the same argument expecting purifying selection at GPAT3, but with ω of 0.25 and little sign of departure from neutral divergence on a codon or species-pair basis, this gene seems to be evolving neutrally. ω for LOG was 0.12, and this gene shows a pattern of divergence quite similar to the nearly neutral behavior of GPAT3. Finally, the analysis of cytidine deaminase needs to be partitioned into the X-linked and Y-linked copies though both gave similar results with ω of 0.16.

### Synteny with Oil Palm

A majority of male-specific date palm BAC contigs had DNA sequence similarity to a single oil palm scaffold (NCBI refseq nucleotide ID NW_011550905.1) (Figure 3a,b) and gene order was strictly maintained between oil and date palm (Figure 3b,c). Date palm BAC contigs with the highest density of male-specific kmers contained *CYP703* and *GPAT3*. These genes are in close proximity in the oil palm locus but missing from the syntenic region in both X and Y date palm alleles. Nevertheless, full-length copies of both genes were detected in all males only (Figure 3d). There was a truncated form of the *GPAT3* gene in the syntenic region with the same retained portion of the 3’ end in both X and Y alleles, indicating that recombination between these alleles continued at this site even after the initial loss of *GPAT3* from the proto-X. Interestingly, the male-specific BAC contig containing the full-length *GPAT3* gene showed similar truncated forms of the gene just downstream of the full-length copy. Based on this observed possible instability in the region, we infer that a duplication occurred prior to the *GPAT3* deletion in proto-X.

Normalization of the sequence read coverage to genome-wide coverage indicated that both *CYP703* and *GPAT3* are maintained as haploid (1N) (Figure 3d) and strengthens the theory that these are indeed unique to the Y chromosome.

For the date palm sequence surrounding the *CYP703* and *GPAT3* deletion in both X and Y alleles, we were able to extend the male allele sequence and noted that the region was fused to a location syntenic to the beginning of the oil palm contig. The fusion point is located between a Myb315-like and a cytidine deaminase-like gene in oil palm with the Y-linked cytidine deaminase in date palm having been moved from the syntenic region (dpBY2) (Figure 3). The Myb315-like gene has already been identified^14^ and studied^18^ in depth as a Y-linked gene in the genus *Phoenix*. Whether this fusion point is present in the female allele is not clear. However, the fact that the cytidine-deaminase contains sequences conserved in all *Phoenix* males investigated suggests that the fusion of the region may have played a role in the formation of the Y chromosome. In addition, an inversion with respect to date palm BAG-like gene was noted in only the male allele (Figure 3c). We identified the Y-linked date palm cytidine deaminase gene in one of the repeat-rich male-specific BAC contigs (dpB3Y) that could not be extended to join any other contigs.

The only male-linked sequence that did not show synteny to the oil palm scaffold was the BAC contig containing the LOG-like gene (Figure 3). The highest similarity date palm sequence to the male-specific *LOG* sequence was an apparent autosomal paralog that was covered by BAC sequencing (Supplementary Figure 4). There was high sequence similarity in the exons between autosomal and Y-linked paralogs, however there were large numbers of male-specific kmers in introns and apparent regulatory regions of the gene making the two loci easily distinguishable (Figure 3d).

### Gene expression analysis

Gene expression analyses of male and female leaves and flowers revealed that expression of *CYP703* and *GPAT3* only occurred in male flowers (Figure 3d). Cytidine deaminase, whether from X or Y alleles, did not show significant expression in any of the four tissues investigated. The *LOG* gene was highly expressed in male flowers with weaker expression in female flowers. However, as mentioned, the exonic region of *LOG* did not contain kmers conserved in all males. SNP analysis showed that the reads likely derived from the autosomal copy of the gene, though the very high levels of expression in the male flower may still be of interest. Expression in male flowers appeared higher in the five most 3’ exons, whereas expression in female flowers was low, but even across the gene body. Whether this was a result of functional differences will require further investigation.

### Discussion

One of the more puzzling aspects of the evolution of sex chromosomes is their origination from ancestral species that lack sex chromosomes. Many plants appear to have relatively new sex chromosomes and therefore offer an opportunity to understand these processes. Localizing the genes critical to sex determination and the steps in sex chromosome evolution is challenging given the suppression of recombination frequently seen between the sex chromosomes and the accompanying nucleotide sequence divergence between the X-specific and Y-specific alleles. Here, utilizing genome sequencing of all members of the genus we identified a small number of genes that exist only in the males of the dioecious *Phoenix*. This is despite the fact that divergence between X and Y has since spread over multiple megabases during speciation^14,15^. All 3 of the most highly conserved male sequences including *CYP703, GPAT3* and *LOG* appear to be maintained in single copy in males. This would be expected if they were deleted from females and only reside on the non-recombining portion of the Y chromosome in males. Our approach requires that divergence between all the species in a genus is sufficiently low that sex-specific genes have enough conserved kmers to be detectable. Indeed, TBLASTN analysis of the male BAC contigs identified sequences such as MAF1 downstream of cytidine deaminase, having sequence similarity to known genes but no male-specific kmers. These genes are likely not critical to dioecy as they have degraded faster among species.

### Analysis of male-specific gene function

Functional studies in other plant species have shown that *CYP703* and *GPAT3* play a critical role in male reproductive organs and male fertility, while *LOG* genes are associated with female flower development. Specifically, mutants of *CYP703* in rice^21^ and maize^29^ and *GPAT3* in rice^24^ revealed their function in both pollen formation and/or anther development through their indispensable role in various lipid synthesis pathways. In rice, *GPAT3* and *CYP703A* are expressed in tapetal cells^24,30^ which are responsible for synthesis and secretion of sporopollenin precursors, a major component of the outer pollen cell wall^31^.

Deletion of *GPAT3* in rice resulted in complete male sterility while normal vegetative development was unaffected. The sterile phenotype was inherited in a recessive fashion with plants carrying a single copy retaining fertility. This strengthens our theory that a single copy Y-linked *GPAT3* in *Phoenix* likely retains sufficient gene function for normal male flower development. In rice *gpat3* mutants, most pollen grains had abnormal pollen wall development, never fully matured and were later aborted^24^, a phenotype that resembled rice *CYP703A* mutants. Of high interest was that the expression of *CYP703A3* was significantly reduced during flower development in rice *gpat3* mutants, suggesting the possible functional connection of these two genes.

The *CYP703* is a single member gene family found across land plant taxa, suggesting it encodes for an essential function^32^. As such, a paralog in *Phoenix* likely does not exist, and deletion of the gene in the proto-X would not be compensated for. Knockout of *CYP703* homologs in Arabidopsis^20^, rice^21^ and maize^29^ resulted in male sterility with no apparent effect on vegetative growth. The *LOG* or LONELY GUY family of genes is involved in activation of cytokinin, an important phytohormone. Functional analysis of the first identified *LOG* member in rice showed that all rice *LOG* mutants were impaired in normal flower development, producing flowers that lacked ovules^33^ or flowers with no pistil and a single stamen^25^. Furthermore, these mutants developed normal inflorescence early on, but then aborted floral meristems. This agrees well with the observations that early flower development in date palm advances both flower organs followed by abortion of one organ depending on the sex of the tree^34^.

The only sequence with conserved kmers in all investigated males that has no clear floral function is the cytidine deaminase-like gene located at the border of the fusion between sequences surrounding the *Myb*315-like and *BAG* genes. Furthermore, X and Y chromosomes maintain a copy of the gene, although it is unclear if both are functional. It is possible that the cytidine deaminase was simply a passenger of the process of sex chromosome evolution due to its proximity to the chromosomal inversion.

### A model for the evolution of dioecy in the genus *Phoenix*

The few conserved male-specific genes absent in all females in the genus simplifies the process of modeling the early progression of sex determination in *Phoenix*. It suggests that speciation in the genus was either concurrent with or immediately followed development of sex determination. The data are also consistent with speciation happening before complete geographical separation occurred. Indeed, hybridization can still occur among different species in the genus and this may also have minimized X and Y divergence between the clades.

The fact that two of the three most conserved male-specific genes derive from a single locus, and the fourth male-linked gene is at the border of a chromosomal rearrangement within the same region make this the most likely candidate region involved in sex determination. The simplest model would suggest that the genus *Phoenix* progressed from hermaphroditism to gynodieocy through the loss of male function by either *CYP703* and/or *GPAT3* deletion on what was the proto-X chromosome. This would leave both genes intact on the proto-Y (Figure 4). The recessivity of these male-sterility mutations agree well with the model proposed by Charlesworth and Charlesworth^2^. The existence of the proto-X would result in gynodioecy – homozygotes of the *CYP703*/*GPAT3* deletion would be female, while heterozygotes and homozygotes carrying the “normal” and ultimately proto-Y allele would remain hermaphrodites. The chromosomal state conferring females would lose male flowering and would spread in the population through the increased efficiency of female seed production^35^. A chromosomal rearrangement that fused the region between the Myb315-like and BAG-like genes then occurred on the “normal” chromosome resulting in recombination arrest between the proto-Y and proto-X. This may have been an inversion that moved the cytidine deaminase, *GPAT3* and *CYP703* genes to a chromosomal end, allowing them to remain on the male allele though not in the original region syntenic with oil palm. The pattern of strong purifying selection on *CYP703* is consistent with this critical male function, although it is less clear why *GPAT3* did not also show this pattern, except possibly that this protein is simply more tolerant of amino acid substitutions. Lastly, the *LOG* gene was translocated into the proto-Y non-recombining region and led to the suppression of female flowers in males, possibly through a dominant mutation that suppresses normal *LOG* function by expression of a truncated LOG competitor. The nearly neutral pattern of divergence of LOG on the Y chromosome might be consistent with this negative function; the constraints on the gene to have female suppressive effects might be considerably relaxed, consistent with its nearly neutral divergence among *Phoenix* species.

While alternative models are possible, this model seems simplest. It agrees well with the theoretical model proposed by Charlesworth and Charlesworth^2^. It is important to stress that the main region of divergence between X and Y is centered on a locus containing the deletion of two genes from females whose deletion in other monocots results in male sterility. The model further predicts the translocation of a female suppressor, a second locus, into the non-recombining region. The *LOG* gene is the only conserved male-specific gene whose family function is known to be important to female flower function and appears to have been duplicated and translocated onto the Y chromosome. Whether *LOG* is indeed a dominant suppressor of female function will require further research. Altogether, our results provide evidence that *Phoenix* passed through a gynodioecious intermediate in development of dioecy and supports a two-locus model of sex determination in the genus. Our results support the theory that dioecy developed prior to speciation in *Phoenix*. Of high interest is that male-specific sequences maintained across the genus include only four genes, three of which are known to be important in male and female flower function in other monocots. Whether these gene sequences could allow for gynodioecy or dioecy to be introduced into close relatives such as oil palm will be of biotechnological interest in the future.

## Methods

### Sample collection and genome sequencing

Leaves of *Phoenix* species were collected by taxonomic and palm specialists in the field at various locations including the USDA in California, the Universidad Miguel Hernandez in Alicante, Spain, and the Huntington Gardens in California (Table 1). Leaves from multiple individuals from all fourteen *Phoenix* species^36^ were collected and one male and one female individual of each species was sequenced and analyzed in this study (with the exception of *P. pusilla* for which a true to type male could not be found). Leaves from the monoecious palm species *Livistona rotundifolia* and *Brahea dulcis* were included for control. DNA was extracted from leaf material using the DNeasy Plant Kit (Qiagen). DNA sequencing was conducted on the Illumina 2500 and 4000 according to the manufacturers recommended paired-end sequencing protocol with libraries of ~400bp. Sequence coverage was determined by analysis of the most prominent kmer peak using JELLYFISH^37^ and by alignment to the date palm genome followed by empirical inspection of coverage at 5,000 SNP sites in 3 scaffolds not showing linkage to sex determination.

### Kmer analysis

We attempted to find kmers of defined sequence length conserved in all members of one sex yet absent in all members of the other. First, to determine the kmer length of appropriate sensitivity and specificity, we identified the maximum length of kmer that was likely to be conserved in all males of the genus despite normal sequence divergence. To that end, sequencing reads from all species were aligned to the date palm reference (NCBI accession ACYX00000000) using BOWTIE2^38^ and SNPs were called using SAMTOOLS^39^ while avoiding annotated repeats and positions with excessive coverage (greater than 3X the mean coverage). Four scaffolds with either no male associated SNPs in more than 6 species were utilized for analysis (dpS5505H2, dpS5560CH1, dpS5784H1, dpS5828CH2) spanning a total of 2,870,261bp. Non-repetitive sequence of 1,780,052bp from the selected scaffolds contained 107,692 SNPs identified in at least one of the 13 male genomes. This revealed a rate of approximately one polymorphism per 16.5 bp among all male members of the species tested. Distances between polymorphisms in the genus were not normally distributed with SNPs more likely to cluster near each other (data not shown). Indeed, 68% of SNPs occurred within 16 bp of each other. However, scanning the test regions in 500 bp windows with a step size of 250 bp we did not identify any of the 11,483 windows that had less than 20 sites (40 including reverse complement) of 16 bp fully conserved across all males. Therefore selecting a kmer size of 16 bp would have an empirical probability of less than 8.7 × 10^-5^ of not identifying a 500 bp or greater fragment of DNA that was unique to males in the genus.

To identify sex-specific kmers, 16 bp kmers were extracted from the raw sequencing read (FASTQ) files using JELLYFISH^37^ for each genome. A genome-specific cutoff was selected based on coverage (ranging from 3–15) under which kmers were assumed to derive from errors in reads and were discarded. Files of male-or female-specific kmers were created for each species by removal of kmers matching those in the opposite sex of that species. Sequencing reads containing kmers present in 13 males and absent from all 14 females were selected from the genome FASTQ file of the Deglet Noor BC5 male. These were assembled to create short contigs on which PCR primers were designed to screen a date palm BAC library. Male-specific sequences were then validated by checking for their presence in an additional 13 males or females from various species of *Phoenix* (Supp Table 1).

### Sequencing, assembly and annotation of male and female alleles

Two BAC libraries (Amplicon Express Inc. Pullman, WA, USA) were constructed from fresh leaves of *Phoenix dactylifera* Deglet Noor BC5 male to a predicted coverage of 6X each for a total of 12X genome. Libraries were constructed in the pCC1 BAC cloning vector with an average insert size of 125 kb using a partial digest of *HindIII* or *EcoRI*.

PCR primers were designed in regions of the date palm genome previously identified as linked to sex^14^ to probe the date palm male BAC library according to the manufacturer’s recommended protocol (Amplicon Express Inc. Pullman, WA, USA)^40^. DNA from BAC clones identified by PCR screening was extracted from overnight *E. coli* cell cultures using the Large-Construct kit (Qiagen). Libraries were constructed and sequenced on the Pacific Bioscience RSII sequencer according to the manufacturer’s recommended protocol for *de novo* sequencing (Pacific Biosciences, Melon Park, CA). Libraries were size-selected from 10 to 40 kb using BluePippin (Sage Science, Beverly, MA). Only reads longer than 10 kb were utilized to assemble individual BACs utilizing the HGAP assembler (Pacific Biosciences, Melon Park, CA). Effectively all BACs assembled into a single contig. Consensus sequences of multiple BACs were obtained using Gap5 v1.2.14^41^.

Sequenced BAC clones were checked for the presence of X-or Y-allele specific polymorphisms. Where possible, a clone from each allele was selected for extension by further BAC library probing and sequencing.

For phased allele sequencing, the Deglet Noor BC5 (DNBC5) genome was subjected to 10X Genomics (Pleasanton, CA) library construction and sequencing per the manufacturer’s protocol. Data was output with the ‘pseudohap2’ model that attempts to separate haplotypes during sequence assembly. Male-specific kmers found in the DNBC5 genome and at least 2 other species’ males but not their counterpart females were used to search the phased haplotype scaffolds for X and Y linked scaffolds. Scaffolds with at least 100 male-specific kmers and 6 times more male-specific kmers in one haplotype were selected for further analysis (Supplementary Table 3).

For annotation, REPEATMODELER was used to identify new repeats followed by REPEATMASKER^42^ to mask repeats in BAC contigs and 10X scaffold sequences. Gene predictions were obtained using a combination of the programs FGENESH++ (Softberry Inc., USA), Augustus^43^, SNAP^44^ and manually curated gene models. The putative role of the candidate genes was determined by BLAST searches against the NCBI database and the oil palm annotated reference genome ASJS00000000^45^.

### RNA-seq and gene expression analysis

Date palm flowers and leaves were collected from two separate male and two separate female trees at anthesis during the flowering season of 2016 in Doha, Qatar. Total RNA was extracted according to the protocol described by Chang and colleagues^46^ followed by on-column DNase treatment using the RNeasy plant mini kit (Qiagen). mRNA-seq libraries were constructed using the Ovation RNA-seq V2 kit (NuGen, San Carlos, CA) according to the manufacturer’s protocol. Libraries were sequenced using either 75bp or 150bp read-lengths on a HiSeq4000 (Illumina, CA) according to the manufacturer’s recommended protocol. Sequences were quality trimmed and aligned to repeat-masked scaffolds using HISAT2^47^. Sequence counts were normalized to fragments mapping per kb of non-sex-linked control scaffold sequence.

### Phylogenetic analysis

Phylogenetic analysis of genes was conducted on *de novo* assembled sequences for each species where possible. Briefly, for each species’ male, genome sequence reads matching the male-specific kmers were collected with their paired-end mates and assembled using the SPADES assembler^48^. Where possible, the same approach was used on the samples from the hermaphroditic genus *Brahea* and *Livistona*. Consensus sequences for each species were then trimmed to allow for global alignments. For cytidine deaminase, male-specific kmers were too few to collect enough sequence for routine assembly. Therefore, the female reference sequence was used and SNP positions substituted with male-specific alleles. Multiple sequence alignments and phylogenies were inferred using MEGA7 with default parameters^49^. Coding sequences were aligned with ClustalW and phylogenies were inferred by Maximum Likelihood using the Jukes-Cantor model based on 1000 bootstraps. Synonymous (dS) and nonsynonymous (dN) substitution rates were calculated based on the Nei-Gojobori method implemented in SNAP v2.1.1^50^. A set of 22 genes conserved in all plant genomes termed “universal single copy orthologs”^26^ were selected from date palm and oil palm for understanding average synonymous substitution rates between the two species.

### Data Availability

All contig/scaffold has been deposited in Genbank under the Accession [HOLDER]. All species’ whole genome sequencing reads have been deposited in the Sequence Read Archive at NCBI under the Accession [HOLDER].

## Acknowledgments

We thank Sean Lahmeyer at the Huntington Gardens for his kind assistance with collection of palms for this study. We thank Encarnacion Carreño from the University of Murcia and Concepcion Obón from the University of Miguel Hernandez (National Phoenix Palm Germplasm Repository of Spain) for their assistance in collection of *Phoenix* species. This study was made possible by grant NPRP-EP X-014–4–001 from the Qatar National Research Fund (a member of Qatar Foundation).

## Author contributions

M.F.T conducted the BAC library probing, RNA-seq libraries, assembly, phylogenetic analysis and wrote the manuscript, L.S.M conducted genome sequencing, library construction and helped write the manuscript, I.A. conducted bioinformatics analysis and helped write the manuscript, I.K.A.A conducted RNA-seq libraries and sequencing, R.K. maintained palm collections, provided phenotyping and systematics analysis, D.R. maintained palm collections, provided phenotyping and systematics analysis, Y.A.M. directed in library construction and sequencing and conducted bioinformatics analysis, A.G.C. conducted evolutionary analysis and helped write the manuscript, K.S. conducted bioinformatics analysis and helped write the manuscript, J.A.M envisioned the project, conducted bioinformatics analysis and wrote the manuscript.

## Materials and Correspondence

Correspondence and BAC requests should be addressed to J.A.M. Palm materials requests should be directed to R.K. and D.R.

